# Genomic Diversity of *Neisseria gonorrhoeae* Isolates in Kenya Revealed by MLST, NG-MAST, and NG-STAR Typing

**DOI:** 10.1101/2025.10.19.683319

**Authors:** Mary Wandia Kivata, Fredrick Lunyagi Eyase, Wallace Dimbuson Bulimo, Valerie Oundo, Esther Waguche, Wilton Mwema Mbinda, Margaret Mbuchi

**Affiliations:** Department of Biological and Physical Sciences, Karatina University (KarU), Karatina, Kenya; Kenya Medical Research Institute (KEMRI), Nairobi, Kenya; Department of Chemistry and Biochemistry, Pwani University (PU), Mombasa, Kenya

**Author notes:** **Corresponding author:** / (MWK). **Alternate corresponding author:** (MWM).

**Keywords:** *Neisseria gonorrhoeae*, antimicrobial resistance, MLST, NG-MAST, NG-STAR, genomic surveillance

## Abstract

**Background:** Surveillance of *Neisseria gonorrhoeae* strains, their antimicrobial resistance (AMR) profiles, and transmission dynamics is essential in the prevention and control of gonococcal infections. In Kenya, gonococcal molecular surveillance remains limited, leaving gaps in understanding circulating sequence types (STs). This study characterized Kenyan *N. gonorrhoeae* isolates using multiple molecular typing schemes.

**Methods:** Illumina MiSeq generated paired-end sequence reads prepared from 35 *N. gonorrhoeae* isolates recovered from males and females from four different regions in Kenya were analyzed. Assemblies were analyzed using PubMLST tools for multi-locus sequence typing (MLST) and the *N. gonorrhoeae* sequence typing for antimicrobial resistance (NG-STAR) scheme. Multi-antigen sequence typing (NG-MAST) was carried out using the NG-MAST database. Phylogenetic relationships were assessed using concatenated NG-STAR loci and core genome-based analyses.

**Results:** Twenty-two MLST STs were identified, including eight novel STs; ST-1932 was most frequent. NG-MAST revealed 29 STs, of which 26 were novel, with a newly described ST-19168 predominating in Nyanza region. NG-STAR identified 23 STs with variation across *mtrR*, *penA*, 23S *rRNA*, *gyrA*, *parC, ponA*, and *porB.* Phylogenetic analyses showed clustering of isolates into distinct groups with diverse AMR profiles. One cluster comprised isolates resistant to tetracycline and ciprofloxacin. No clear association was observed between MLST, NG-MAST, or NG-STAR types and specific AMR patterns.

**Conclusion:** Kenyan *N. gonorrhoeae* strains are genetically diverse, with high numbers of novel NG-MAST and MLST STs. The lack of regional clustering and varied AMR profiles suggest widespread transmission of heterogeneous gonococcal populations. These findings underscore the importance of strengthened genomic surveillance to inform gonorrhea control strategies in Kenya.

## Background

Management of gonococcal infections is complicated by the emergence and spread of *N. gonorrhoeae* strains which are resistant to the current antibiotics recommended for treatment [1]. In addition, prevention and control of infections is hindered by the lack of a gonococcal vaccine[2]. Identification and genotypic characterization of these antibiotic resistant strains through molecular typing is a key factor in monitoring the emergence, transmission and spread of gonococcal drug resistance [3]. Characterization of sequence types and the associated antibiotic susceptibility profiles of circulating *N. gonorrhoeae* isolates provide epidemiological and drug resistance data useful for management and prevention of gonococcal infections including, updating of treatment guidelines and other preventive interventions.

Both DNA-based (genotypic) and non-DNA-based (phenotypic) typing methods have been developed and used to characterize gonococcal isolates. Phenotypic typing methods include: antimicrobial susceptibility testing (AST); auxotyping; and serotyping[3]. They are limited in terms of reproducibility and their ability to discriminate between strains. DNA-based typing methods are categorized into gel-based and sequence-based typing methods. Gel based typing methods employ gel electrophoresis to analyze DNA bands and include: ribotyping; restriction fragment length polymorphism (RFLP) and pulsed-field gel electrophoresis (PFGE); opa-typing; and PCR based typing[3]. DNA sequence-based typing methods entail the analysis of one or more genes and include; *Neisseria gonorrhoeae* Multi-antigen Sequence Typing (NG-MAST); Multi-locus Sequence Typing (MLST); and a recently described scheme called *N. gonorrhoeae* Sequence Typing for Antimicrobial Resistance (NG-STAR)[4, 5]. DNA sequence-based typing methods are preferred because: they have high discrimination power; are rapid and reproducible; allow identification of novel polymorphisms and also transfer and comparison of results [3].

MLST uses partial sequence information of seven relatively conserved, slow evolving housekeeping genes which are distributed throughout the genome (*abcZ*, *adk*, *aroE*, *fumC*, *gdh*, *pdhC*, and *pgm*) to assign an isolate to a sequence type [5, 6]. It is recommended for typing gonococcal infections in long-term and global epidemiology studies [3]. NG-MAST analyses partial DNA sequences from hypervariable regions of two outer membrane proteins; porinB encoded by *porB* (490pb) and, transferrin binding protein encoded by *tbpB* (390bp). It uses an open access database (http://www.ng-mast.net/) for analysis and is recommended for typing gonococcal infections in micro-epidemiological studies [3, 4]. NG STAR is a newly described antimicrobial resistance multi-locus typing scheme which is based on gonococcal antimicrobial resistance determinants. It uses alleles from seven genes associated with resistance to β-lactams, macrolides, and fluoroquinolones antibiotics. The genes include: *penA*; *mtrR*; *porB*; *ponA*; *gyrA*; *parC*; and 23S rRNA. This typing scheme provides insights on chromosomal antimicrobial determinants and is useful in monitoring global dissemination of antimicrobial resistant gonococcal strains [7].

Gonococcal antimicrobial susceptibility testing combined with DNA sequence-based typing data provide information useful in: understanding strains evolution and genetic relatedness; monitoring changes in existing strains and emergence of new drug resistant strains; and monitoring the spread of antibiotic resistant strains in specific groups or populations [3, 8]. In Kenya few studies have characterized *N. gonorrhoeae* whole genome sequence data [9, 10]. Consequently, data on the current circulating gonococcal sequence types is limited. This study used the commonly used and recommended DNA sequence-based typing methods to characterize genetic heterogeneity of *N. gonorrhoeae* isolates obtained from different regions in Kenya.

## Methods

### Study isolates

Study isolates consisted of 35 *N. gonorrhoeae* isolates obtained from both male and female patients seeking treatment in selected clinics from four different regions in Kenya: Nairobi; Coastal Kenya; Nyanza and Rift Valley between 2013 and 2018. The isolates were collected and antimicrobial susceptibility testing done as part of an ongoing STI surveillance study titled “A surveillance study of antimicrobial susceptibility profiles of *N. gonorrhoeae* isolates from patients seeking treatment in selected clinics in Kenya”. Minimum inhibitory concentrations for: ceftriaxone; cefixime; penicillin; tetracycline; azithromycin; ciprofloxacin and spectinomycin; (S1 Table) [11] were determined using E-test® (Biomerieux) method according to manufacturer’s instructions [12, 13] and the breakpoints interpreted with reference to European Committee on Antimicrobial Susceptibility Testing (EUCAST) version 8.0, 2018 standards (S2 Table). DNA isolation and whole genome sequencing was done as part of a retrospective laboratory based molecular sub-study titled “Molecular characterization of antimicrobial resistance genes in *Neisseria gonorrhoeae* isolates from Kenya through whole genome sequencing” which was nested in the STI surveillance programme. Genomic DNA was extracted using QIAamp DNA Mini Kit (QIAGEN, Hilden, Germany) and the quality and quantity of DNA determined by Qubit dsDNA HS Assay using Qubit 3.0 fluorometer, (Thermo Fisher Scientific Inc. Wilmington, Delaware USA) according to the manufacturer’s instructions.

### Whole Genome Sequence Data, Assembly and Annotation

Illumina Nextera XT kit (Illumina Inc. San Diego, CA, USA) was used to prepare libraries as per manufacturer’s instructions. Sequence reads were generated on Illumina MiSeq platform (Illumina, San Diego, CA, USA) using a paired-end 2×300bp protocol[14]. Sequence reads were assembled with CLC Genomics Workbench (v12.0, Qiagen). Genome annotation and analysis were conducted with BIGSdb tools accessed through the PubMLST platform (https://www.pubmlst.org/neisseria). The generated sequence reads are linked to NCBI BioProjects: PRJNA481622 and PRJNA590515 and the assemblies are available on PubMLST.

### Multi-locus Sequence Typing, NG-MAST and NG STAR

Identification of multi-locus sequence types (MLST) was performed on assembled genome sequences using MLST version 1.8 (https://cge.cbs.dtu.dk/services/MLST/) available online at Centre for Genome Epidemiology (CGE) (https://cge.cbs.dtu.dk/services) [15]. A local blast search against NG-MAST *porB* and *tbpB* database was used to identify *porB* and *tbpB* genes from the assembled genomes using Bioedit sequence alignment editor version 7.0.5 [16]. Determination of *porB* and *tbpB* alleles and NG-MAST sequence types (STs) was done at NG-MAST website (http://www.ng-mast.net/) using correctly trimmed *porB* (490bp) and *tbpB* (390bp) genes. Determination of *N. gonorrhoeae* Sequence Typing for Antimicrobial Resistance (NG STAR) STs was done using the NG STAR scheme implemented in PubMLST [7].

### Phylogeny analysis

To identify phylogenetic relationships among the study sequences, genome comparator tool hosted on https://pubmlst.org/neisseria was used to compare whole genome sequence and core genome-based phylogeny inferred using cgMLST *N. gonorrhoeae* v.1.0 scheme. The generated data was visualized using Inkscape. ITOL tool hosted on https://pubmlst.org/neisseria was used to create a neighbor-joining tree from concatenated seven NG-STAR loci nucleotide sequences. The core genome comparison data was annotated by NG-STAR, NG-MAST and MLST schemes. To analyze clustering of sequence types in global context, ITOL was used to generate a neighbor-joining tree based on NG-STAR, NG-MAST and MLST schemes. Thirty-nine additional global strain sequences from different countries were acquired from PubMLST and included in the analysis for comparison purposes.

### Statistical Analysis

All proportions are reported with 95% confidence intervals (CIs) using the binomial exact method. Statistical comparisons were performed using Mann–Whitney U tests for continuous variables and Fisher’s exact tests for categorical variables. Given the small sample size and exploratory nature of this study, p-values were not corrected for multiple testing and should be interpreted with caution. We abstracted sex, age, marital status, and self-reported partner count from clinic records (S1 Table). For partner count, categories were collapsed to >2 vs ≤2/none due to sparse upper cells. Genomic novelty was indicated by the presence of a novel sequence type in any scheme (MLST, NG-MAST, NG-STAR). For typing-AMR analyses, frequent sequence types (MLST ST-1932, ST-8133; NG-MAST ST-19168; NG-STAR ST-1586, ST-2668) were compared vs all other types for each drug. Given small sample sizes and sparse cells, results are interpreted as exploratory.

### Ethical Consideration

Permission to carry out the study for both the STI surveillance study and the nested molecular sub-study were obtained from Kenya Medical Research Institute (KEMRI) Scientific and Ethics Review Unit) and Walter Reed Army Institute of Research Institutional Review Board as (WRAIR#1743, KEMRI#1908) and (WRAIR#1743A, KEMRI#3385) “respectively”. Consent to participate was not applicable for this study because it was retrospective laboratory based, used archived samples, and there was no interaction with human subjects. All data were fully anonymized before we accessed them and the research was qualified as not involving human subjects. The isolates were acquired in two batches on 23/1/2017 and 26/2/2018.

## Results

All the thirty-five isolates were included across all three typing schemes. The isolate IDs are sequential, but not all numbers are represented, reflecting availability of quality-assured WGS data. No isolates were excluded after typing.

### Sequence typing

#### *N. gonorrhoeae* Multi-locus Sequence typing

A total of 22 MLST STs representing: 3 *abcZ*; 2 *adk*; 3 *aroE;* 6 *fumC*; 5 *gdh*; 3 *pdhC;* and 2 *pgm* different alleles were identified. Of the 35 sequences, 24 belonged to 14 known MLST STs; ST-1588; ST-1599; ST-1893; ST-1921; ST-1928; ST-1932; ST-8111; ST-8133; ST-11242; ST-11365; ST-11366; ST-11367; ST-11750; and ST-11976; while the remaining 11 sequences belonged to 8 new STs; ST-13613; ST-13614; ST-13763; ST-13764; ST-13766; ST-13779; ST-13780; and ST-13782 (S3 Table). The predominant ST was ST-1932 (*n* = 5, 14.3%) followed by ST-8133 (*n* = 4, 11.4%) and a novel ST-13782 (*n* = 3, 8.6%). In 2015, 44.4% (95% CI: 18.9–73.3%) of isolates displayed novel MLST types. Similarly, ciprofloxacin resistance was observed in 97.1% (95% CI: 83.4–99.9%) of isolates, underscoring the near-fixation of resistance in the population. Twenty isolates formed seven groups (two or more same ST) while 15 isolates formed singular MLST STs. Unlike ST-1932 and ST-13782, ST-8133 consisted of isolates recovered from one region; Nyanza. Region-based occurrence was also observed in ST-11365 and ST-13780 which consisted of two isolates each from Nyanza (S3 Table).

#### *N. gonorrhoeae* Multi-antigen Sequence Genotyping

In total, 29 NG-MAST sequence types were detected, encompassing 22 distinct *porB* alleles and 21 *tbpB* alleles. Nine of both the 22 *porB* and the 21 *tbpB* alleles were novel. Thirty-one (88.6%) isolates belonged to 26 novel NG-MAST STs: ST-18599; ST-19087; ST-19166-ST-19170; ST-19254-ST-19272 while four (11.4%) isolates belonged to 3 known STs: ST-355; ST-10134; and ST-11752. A novel ST, ST-19168 comprised four isolates from Nyanza was the most common (11.4%). These four isolates belonged to an existing MLST ST-8133 which was the second common MLST. Sequence types ST-19255, ST-19262, and ST-10134 which consisted of two isolates each, were the second most predominant. Four ST groups were formed by 10 isolates, while the rest formed singular NG-MAST STs (S4 Table). NG-MAST based regional occurrence was only observed in two novel NG-MASTs; ST-19168 and ST-19255 which all comprised isolates recovered from Nyanza between 2014 and 2018.

#### *N. gonorrhoeae* Sequence Typing for Antimicrobial Resistance (NG STAR)

Twenty-three NG-STAR STs representing 9 *mtrR*, 5 *penA*, 1 23S rRNA, 3 *gyrA*, 6 *parC*, 2 *ponA*, and 6 *porB* different alleles were identified. Seven (20%) isolates belonged to 6 novel NG-STAR STs: ST-3179; ST-3182; ST-3183; ST-3184; ST-3185 and ST-3186 while the rest of the isolates belonged to 17 known NG STAR STs. The most common STs were ST-1586 and ST-2668 which both comprised 4 (11.4%) isolates each. One of the two predominant NG-STAR ST, ST-2668 comprised isolates from Nyanza which also belonged to the most common NG-MAST, ST-19168 and to the second common MLST, ST-8133. Similar region, NG-STAR, NG-MAST and MLST occurrence was observed in two isolates; KNY_NGAMR 20 and KNY-NGAMR 23 from Nyanza. Nineteen isolates formed 7 NG STAR ST groups while the rest formed singular NG-STAR STs (S5 Table)

#### Temporal Distribution of Novel Sequence Types

Across 35 isolates analyzed between 2013 and 2018, the emergence of novel sequence types varied by scheme and year. Yearly isolate distribution was: 2013 (n=2), 2014 (n=4), 2015 (n=9), 2016 (n=9), 2017 (n=7), and 2018 (n=4). Novel MLST STs were first detected in 2014 and peaked in 2015–2016 (44.4% each), persisting through 2017 (42.9%) but absent in 2013 and 2018. NG-MAST showed the highest rate of novelty overall, with at least one novel ST detected each year and peaks in 2014 (100%), 2015 (88.9%), and 2017 (85.7%). NG-STAR displayed fewer novel profiles, appearing sporadically but rising substantially in 2017 (57.1%) and 2018 (75%) (Fig 1).

**Fig 1.**
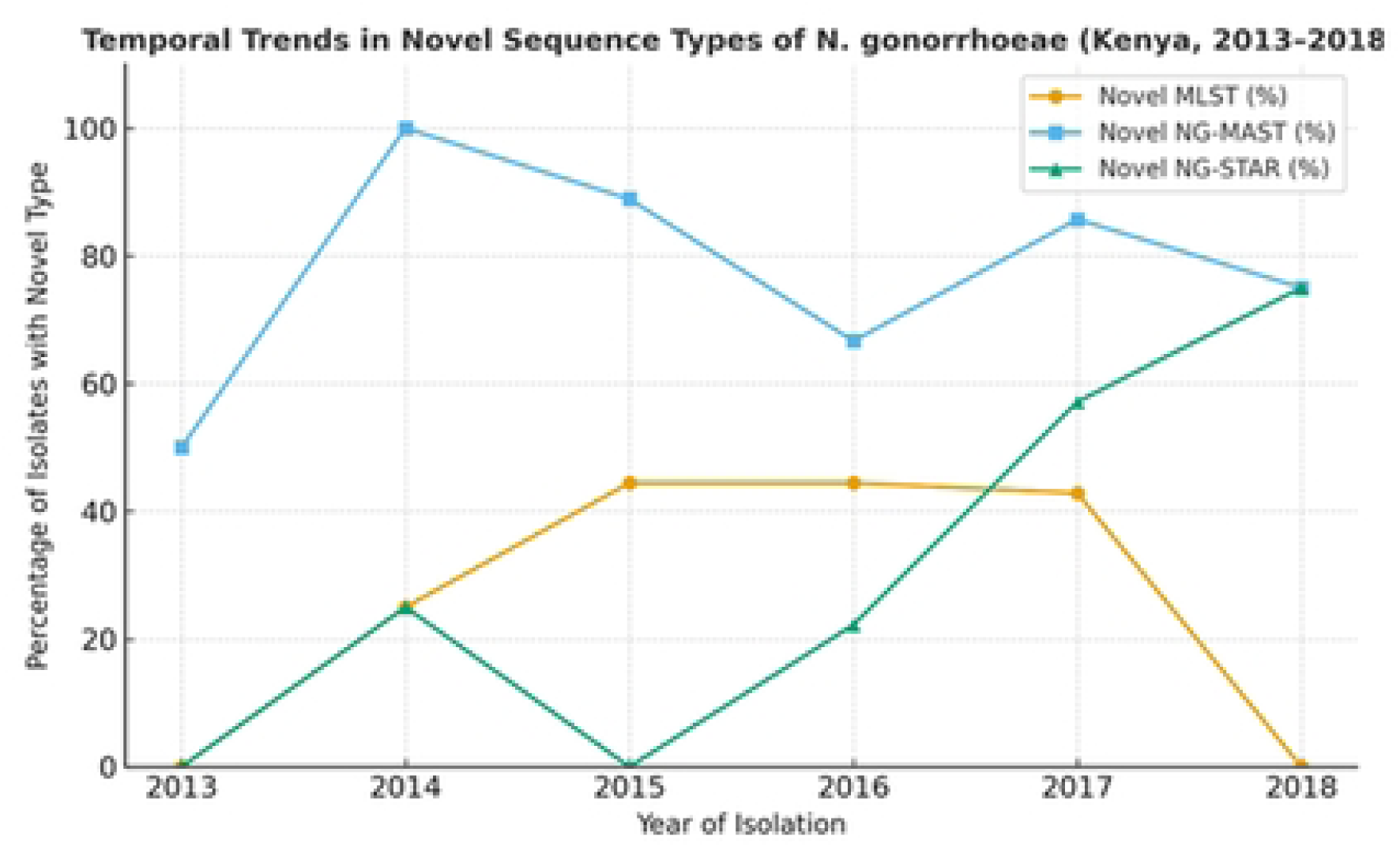
Temporal trends for novel MLST, NG-MAST, and NG-STAR sequence types (2013–2018). Overall, 2015–2017 represented a period of marked diversification across all three schemes, while isolates from 2018 showed stabilization into a smaller number of lineages, albeit still including novel NG-MAST and NG-STAR STs

#### Association between typing schemes and antimicrobial resistance

Resistance to ciprofloxacin, penicillin, and tetracycline was widespread across sequence types (MLST, NG-MAST, NG-STAR). When frequent types were compared to all others, no statistically significant enrichment in resistance was observed for any drug (two-sided Fisher tests, all p≥0.20 (S6 Table). Azithromycin resistance was rare; the non-significant elevation in the NG-STAR ST-2668 group (OR≈4.83; p=0.31) was based on small counts and should be interpreted with caution. Reduced susceptibility to ceftriaxone/cefixime was uncommon and not concentrated in particular sequence types at detectable levels

#### Associations between epidemiological covariates and genomic novelty

Of 35 cases (male 85.7%; median age 26 years, IQR 22–31), “any novel sequence type” was common across demographic strata. No significant associations were detected between novelty and age (median 26.0 vs 30.5 years; U=18.0, p=0.322), sex (male vs female: OR=7.25, p=0.269), marital status (married vs single: OR≈0, p=0.202; sparse zero cell), or partner count >2 (vs ≤2/none: OR→∞, p=1.00; sparse upper category). The high prevalence of novelty across strata suggests broad dissemination of novel lineages in the catchment population rather than concentration in specific demographic groups

### Phylogenetic analyses

#### Core genome-based phylogeny (cgMLST *N. gonorrhoeae* v.1.0 scheme)

Genome comparator tool identified 1485 loci as core genomes to *N. gonorrhoeae*. cgMLST *N. gonorrhoeae* v.1.0 scheme identified four clusters (group of more than 3 isolates) among the study isolates: Cluster 1 (n= 15); Cluster 2 (n= 6); Cluster 3 (n= 5) and Cluster 4 (n= 4). Cluster 1 was further sub-divided into two smaller but distinct sub-clusters; sub-cluster 1a (n=9) and sub-cluster 1b (n=6) (Fig 2). Isolates in sub cluster 1b were from 3 regions and belonged to 3 different MLST, NG-MAST, and NG-STAR STs. Isolates in sub-cluster 1a belonged to: 5 MLST ST (3 novel and 2 existing STs), 8 NG_MAST STs (6 novel and 2 existing STs) and 4 NG-STAR STs. Eight of the nine isolates in this sub-cluster were all from Nyanza. Cluster 2 was formed by isolates from Nairobi and Nyanza belonging to: 2 existing MLSTs STs; 5 NG-MAST STs and 6 NG-STAR STs. Cluster 3 was formed by four isolates from Nyanza and one from Rift Valley. They belonged to; four different MLST and NG-MAST STs (all novel) and 2 NG-STAR STs. Isolates in cluster 4 were not as closely clustered as in the other three clusters. They belonged to: 3 existing MLST STs; 4 new NG-MAST STs and 4 NG-STAR STs (Fig 2).

**Fig 2.**
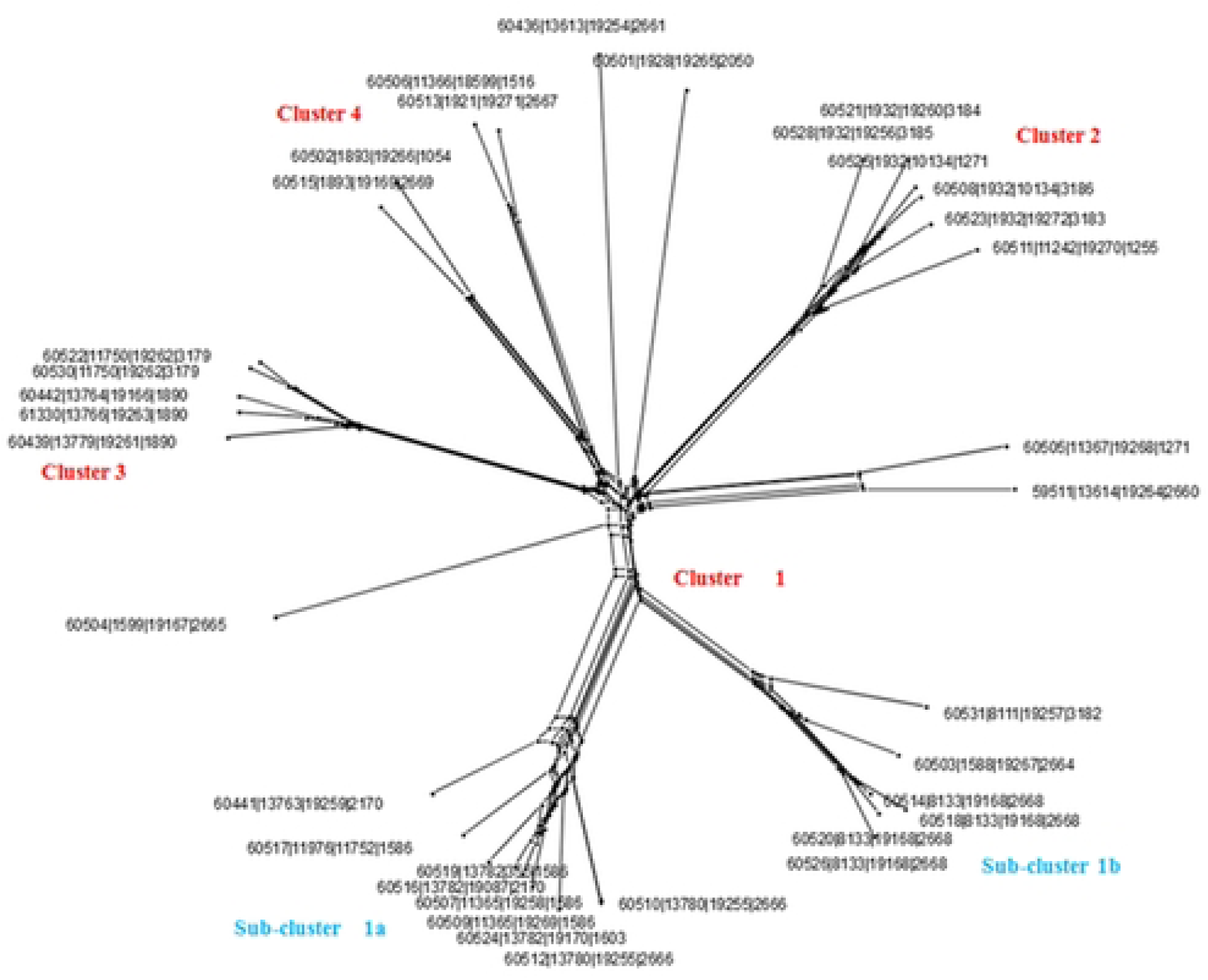
Core genome-based phylogeny inferred from 1485 loci identified as core in 35 Kenyan gonococcal isolates using cgMLST. *N. gonorrhoeae* v.1.0. Four distinct clusters (1-4) were formed. Shown in the leaf terminals are: PubMLST IDs for the isolates; MLST; NG-MAST STs and NG-STAR ST respectively.

#### NG-STAR phylogeny

Neighbor-joining tree inferred by ITOL identified 3 distinct clusters: Cluster A (n= 12); Cluster B (n= 9); and Cluster C (n= 6) (Fig 3). Cluster A was formed by 12 isolates from all the four sampled regions. These isolates had varied antibiotic susceptibility profiles. Ten of these 12 isolates belonged to 6 already known MLST ST. while the remaining 2 isolated belonged to a novel MLST ST. On the contrary more diversity was observed in this cluster with regard to NG-MAST and NG STAR STs as these isolates belonged to ten NG-MAST and ten NG-STAR STs. Nine of these ten NG-MAST STs formed by 10 isolates were novel while one ST formed by two isolates was an existing one (ST-10134).

**Fig 3.**
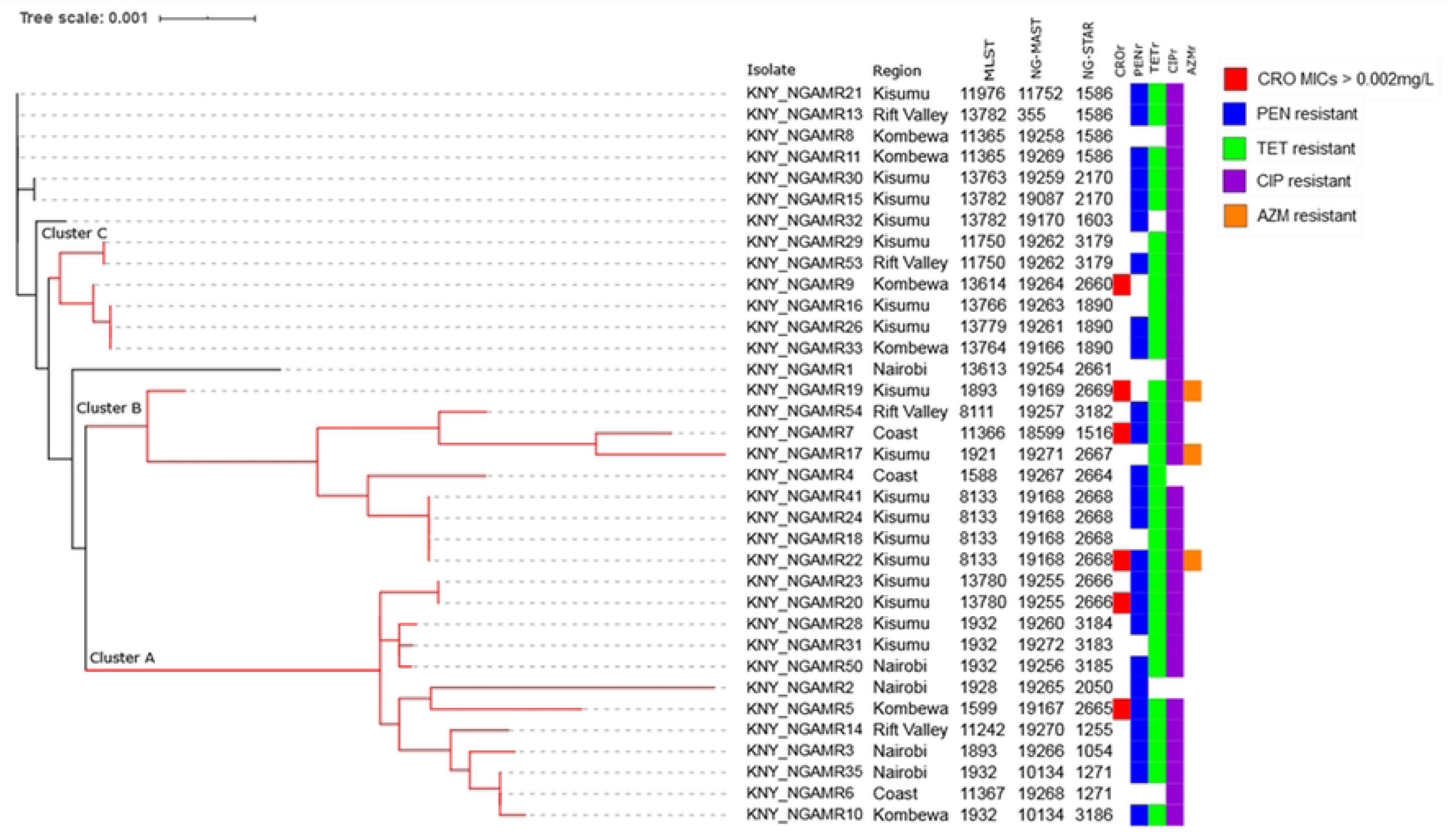
Neighbor-joining tree generated by ITOL from seven concatenated NG-STAR loci nucleotide sequences (n=35) (*mtrR*, *penA*, 23S rRNA, *gyrA*, *parC*, *ponA*, and *porB*). Antimicrobial susceptibility profiles for isolates with reduced susceptibility to ceftriaxone (CRO); (MICs >0.002mg/L) and those resistant to: penicillin (PEN); tetracycline (TET); ciprofloxaxin (CIP) and azithromycin (AZM) have been included in the phylogeny. Three distinct clusters (A-C) formed comprised isolates with varied MLST, NG-MAST and NG-STAR STs. Branch lengths represent evolutionary change or genetic distance between sequences or taxa.

The nine isolates forming cluster B were also from all the sampled regions and belonged to six existing MLST STs. The isolates belonged to six novel NG-MAST STs and six NG-STAR STs. They were all tetracycline resistant, while 8 of the 9 were ciprofloxacin resistant. Additionally, three isolates with a low-level resistance to azithromycin all belonged to this cluster. Cluster C was formed by six isolates; five from Nyanza and one from Rift Valley. Unlike cluster A and B which mostly consisted of existing MLST STs, cluster C isolates belonged to four novel MLST STs and one existing MLST ST. The isolates were all tetracycline and ciprofloxacin resistant (Fig 3) NG-STAR, NG-MAST and MLST region-based clustering was not observed in either of the two phylogenies as the identified clusters comprised isolates with varied STs.

#### Clustering of Novel STs in Global Context

Neighbor-joining tree inferred by ITOL clustered the Kenyan isolates into multiple, distinct and individual clusters. Although genetic diversity was observed in the Kenyan isolates, when compared to global strains they were found to be closely related. Comparative assessment against representative global *N. gonorrhoeae* genomes from PubMLST revealed that most novel Kenyan MLST, NG-MAST, and NG-STAR sequence types did not cluster closely with globally recognized high-risk resistant lineages such as MLST ST-1901 or NG-MAST ST-1407 (Fig 4). African sequences especially those from Uganda, Cameroon and Malawi clustered near Kenyan sequences. Most of these isolates belonged to MLST ST-7363 which is globally associated with ciprofloxacin, penicillin, tetracycline resistance in gonococci [17].

**Fig 4.**
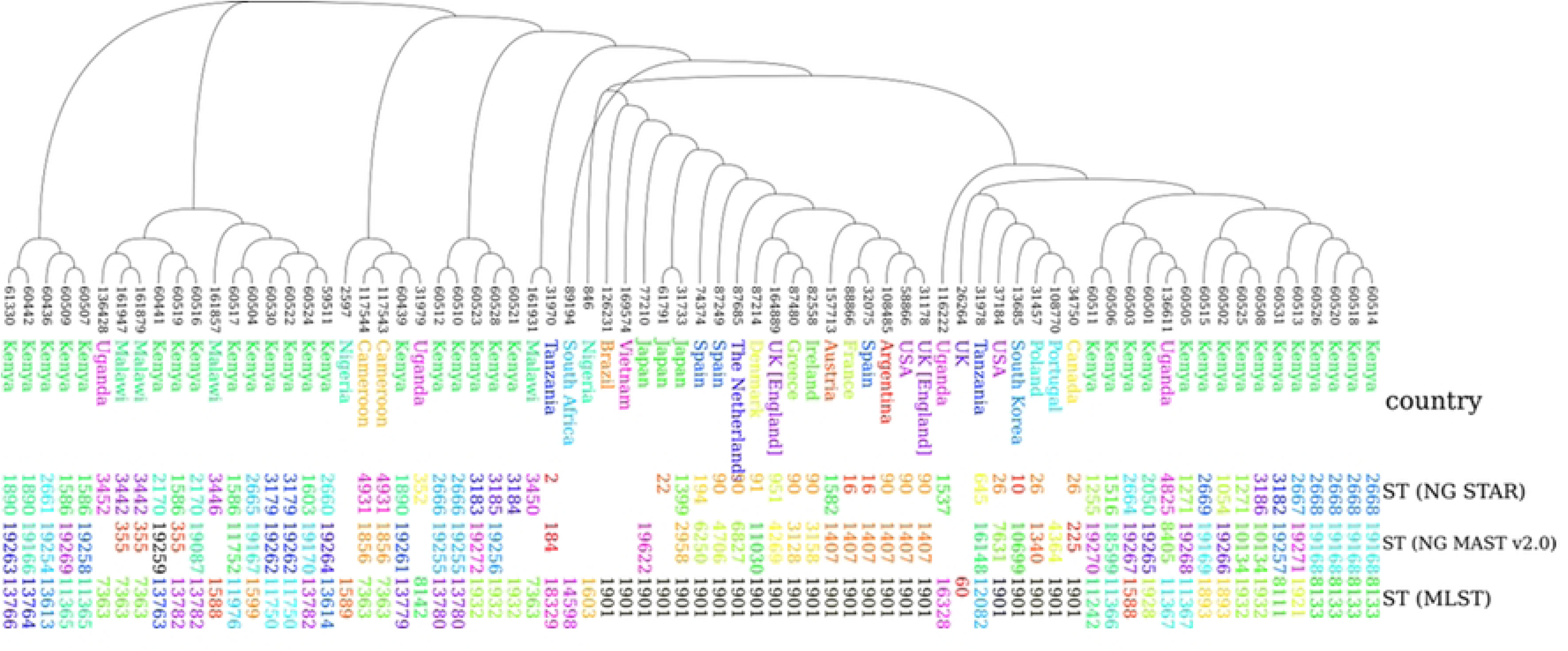
Neighbor-joining tree generated by ITOL using MLST, NG-MAST and NG-STAR schemes (n=74). The Kenya sequence types did not cluster closely with globally recognized high-risk resistant lineages such as MLST ST-1901 or NG-MAST ST-1407. African sequences clustered near Kenyan sequences. The other global strains were included for comparisons. PubMLST IDs for the isolates are shown in the leaf terminals.

## Discussion

Varied sequence types indicate genetic diversity in bacterial isolates. Although the present study identified variety of existing and known MLST, NG-MAST and NG-STAR STs, most of the identified NG-MAST and NG-STAR STs were new to the PubMLST database and were described for the first time in the present study. Additionally, most of the isolates formed singular STs indicating genetic diversity in the analyzed isolates. These findings indicate independent evolution in the Kenyan gonococci. The most sequence diversity was observed in NG-MAST with 25 isolates belonging to singular NG-MAST STs. Similar sequence diversity was observed in gonococci from Western Australian [18]. Isolates belonging to already known MLST STs were typed into different and varied NG-MAST and NG-STAR STs which indicate that the latter typing methods have a higher discriminatory power when compared with MLST.

The temporal analysis highlights a dynamic gonococcal population structure in Kenya. The mid-study years (2015–2017) were characterized by high levels of novelty across MLST, NG-MAST, and NG-STAR, suggesting simultaneous local diversification and external lineage introduction. This coincides with regional reports of increased genetic diversity in *N. gonorrhoeae* during the same period in South Africa attributed to international connectivity and high recombination rates [19, 20]. By contrast, 2018 isolates displayed a stabilization trend, with no novel MLST types and lower absolute numbers of novel NG-MAST STs. The absence of novel MLST STs among 2018 isolates should be interpreted with caution, as the sample size for that year was small (n=4). Whether this reflects true stabilization of lineages or sampling limitations remains uncertain. However, the presence of novel NG-STAR STs in 2017–2018 indicates ongoing evolution of resistance determinants, underscoring the risk of emergent AMR phenotypes even when overall genetic novelty declines. These findings emphasize the importance of continuous genomic surveillance. Without sustained monitoring, early warning signals of resistant lineages may be missed during phases of apparent clonal stability. The Kenyan data thus support an integrated AMR-genomics surveillance framework to capture both rapid diversification events and long-term lineage persistence.

Findings from this study indicate that (i) genomic novelty is broadly distributed across routine demographic strata in the study population, and (ii) legacy resistance markers (CIP, PEN, TET) are diffuse across multiple genetic backgrounds, consistent with long-standing circulation and horizontal gene transfer in *N. gonorrhoeae*. Statistically robust clustering of non-susceptibility within frequent MLST, NG-MAST, or NG-STAR types were not detected in this study. This likely reflects a combination of high background prevalence for legacy resistance, low event counts for macrolide/extended-spectrum cephalosporin resistance, and limited power. From a surveillance perspective, these results argue for continued genomic/AMR monitoring that integrates minimal structured metadata (such as partner concurrency and travel) to resolve transmission micro-structures that routine demographics cannot capture.

Generally, region-based clustering was not observed in present study since the clusters generated from both neighbor-joining and core genome-based phylogenies comprised isolates from varied regions. Although the study sample size is small, these observations suggest that the Kenyan gonococci sampled belong to heterogeneous population. Nevertheless, ST based clustering was observed in three: MLST, (ST-11750, ST-13780 and ST-8133), NG-MAST (ST-19262, ST-19255 and ST-19168) and NG-STAR (ST-3179, ST-2666 and ST-2668). Diversity was not only observed in sequence typing but also in the antimicrobial resistance profiles. Penicillin, tetracycline and ciprofloxacin resistance was observed in all the three clusters formed by phylogeny generated from seven loci associated with gonococcal antimicrobial resistance.

Additionally, no particular ST was associated with specific AMR pattern. This shows that resistance to these three antibiotics is widespread in all the sampled regions. Three isolates from Nyanza exhibiting low level Azithromycin resistance belonged to distinct sequence types including already known MLSTs as well as novel NG-MAST and NG-STAR STs.

Marked incongruences were observed when comparing the core genome phylogeny with the NG-STAR phylogeny. These discrepancies highlight differences in evolutionary signals captured by genome-wide versus resistance gene-based analyses. While the cgMLST tree reflects long-term evolutionary relationships across 1,485 conserved loci, the NG-STAR tree focuses on only seven loci directly associated with antimicrobial resistance. These loci are subject to strong positive selection due to antibiotic use, and *N. gonorrhoeae’s* high capacity for horizontal gene transfer allows these AMR determinants to spread rapidly across unrelated genetic backgrounds. Consequently, genetically distant isolates in the cgMLST tree may cluster together in the NG-STAR phylogeny if they share the same resistance alleles, reflecting convergent evolution under drug pressure rather than shared ancestry. Furthermore, recombination events and mosaic gene structures in loci such as *penA* can introduce resistance-conferring alleles into multiple lineages, producing identical NG-STAR profiles in otherwise distinct strains. The reverse scenario is also possible: closely related isolates in the cgMLST tree may display divergent NG-STAR profiles if they have acquired different resistance alleles through recent recombination or if selection pressures have driven allele replacement. These findings emphasize that NG-STAR phylogenies primarily capture recent adaptive events in AMR gene evolution, while core genome phylogenies retain broader evolutionary signals. Therefore, integrating both approaches provides a more comprehensive view of gonococcal population structure and resistance evolution.

Comparative assessment against representative global *N. gonorrhoeae* genomes from PubMLST revealed that most novel Kenyan MLST, NG-MAST, and NG-STAR sequence types did not cluster closely with globally recognized high-risk resistant lineages such as MLST ST-1901 or NG-MAST ST-1407, which are strongly associated with multidrug resistance and have caused multiple treatment failures in Europe, North America, and parts of Asia [21, 22]. Instead, the Kenyan novel types formed distinct branches within the phylogenetic trees, suggesting they represent locally evolved or region-specific lineages rather than recent introductions of pandemic-resistant clones. This observation aligns with the lack of ceftriaxone-or high-level azithromycin-resistant isolates in this dataset. Ugandan and Malawian sequences clustered near Kenyan ones, probably indicating regional relatedness or shared transmission pathways.

A minority of novel sequence types showed loose phylogenetic proximity to internationally disseminated ciprofloxacin-resistant lineages (MLST ST-7363)[17], indicating possible historic introductions followed by diversification under local selective pressures. The predominance of unique branches for Kenyan novel sequence types underscores the potential for endemic evolution in the absence of sustained international strain replacement, but it also highlights the need for continuous genomic surveillance to detect any incursion of globally dominant high-resistance clones.

Although the present study provides valuable insights into the molecular epidemiology of *N. gonorrhoeae* in Kenya, the relatively small sample size (n = 35) collected over a six-year period limits the statistical power and generalizability of the findings. A small dataset reduces the precision of prevalence estimates for specific sequence types (STs) and antimicrobial resistance (AMR) phenotypes, increasing the likelihood of both Type I errors (false associations) and Type II errors (failure to detect true associations). Consequently, rare but epidemiologically important sequence types or resistance profiles may not have been captured, and the observed proportions of novel STs may be over- or under-estimated due to sampling variability. Metadata were restricted to basic demographics, with under-representation of females (14%) and sparse reporting of high-partner categories. These limitations reduced power to detect epidemiological correlates of genomic novelty. Future surveillance should incorporate enhanced metadata, including sexual network information and travel history, to better capture transmission dynamics

The temporal and geographic representativeness is also constrained. Although isolates were obtained from four regions, the sampling frame was not uniform across all areas and years, and the absence of certain high-risk clones such as MLST ST-1901 and NG-MAST ST-1407 cannot be interpreted as definitive evidence of their absence from the Kenyan population. In molecular surveillance, robust detection of low-prevalence but clinically significant genotypes often require hundreds of isolates per year across sentinel sites to achieve high sensitivity and confidence intervals narrow enough for public health decision-making. Future surveillance efforts should employ systematic, continuous sampling across all high-burden regions, with annual isolate counts sufficient to detect STs with a prevalence as low as 1–2% with ≥80% power. This will enable more reliable temporal trend analyses, support phylogeographic inferences, and improve the ability to detect emerging AMR-associated clones in real time.

Kenya’s current gonococcal isolates lack globally dominant epidemic clones such as ST-1901/1407, which are strongly linked to extended-spectrum cephalosporin resistance. Nonetheless, regional and international travel could facilitate rapid importation. We recommend targeted sentinel surveillance among border towns, mobile populations, and key populations (MSM and sex workers) with integration into WHO GASP and GLASS platforms and alignment with the Kenya AMR Action Plan.

## Conclusions and recommendations

This study provides important insights into the molecular epidemiology of *N. gonorrhoeae* circulating in Kenya. The findings reveal that circulating isolates are genetically diverse, with most of the identified NG-MAST and NG-STAR sequence types being novel and reported for the first time. Importantly, no specific sequence type was consistently associated with particular antimicrobial resistance (AMR) patterns, indicating that resistance is widely distributed across different genetic backgrounds. These results suggest a heterogeneous gonococcal population in Kenya, with broad dissemination of both existing and novel resistant lineages. Although limited by the small sample size, this study underscores the need for strengthened continuous, large-scale surveillance to detect emerging resistance determinants and monitor transmission dynamics. The absence of globally dominant high-risk clones such as MLST ST-1901 and NG-MAST ST-1407 offers a window of opportunity to strengthen local containment before such lineages are introduced.

Based on the observed results, the following recommendations are proposed: (i) strengthen gonococcal genomic surveillance and integrate genomic data into the existing national Antimicrobial Resistance (AMR) action plan to enable early detection of resistant lineages, track transmission patterns, and guide timely public health interventions, (ii) establish sentinel surveillance sites and ensure continuous, systematic sampling to capture both emerging and persistent gonococcal lineages, (iii) foster collaboration between microbiology laboratories, public health institutions, and clinicians to ensure timely feedback of resistance data into patient management. Aligning these actions with Kenya’s national AMR action plan will be critical to prevent the spread of highly resistant gonococcal strains and safeguard the effectiveness of current treatment regimens.

## Supporting information

**S1 Table.** Isolate metadata and antimicrobial susceptibility data.

|  | Isolate details |  |  | Metadata |  |  |  | MICs (mg/L) |  |  |  |  |  |  |
| --- | --- | --- | --- | --- | --- | --- | --- | --- | --- | --- | --- | --- | --- | --- |
|  | Sample ID | Year of isolation | Region | Sex | Age | Marital status | No. of sexual partners | CFX | CRO | PEN | TET | CIP | AZM | SPT |
| 1 | KNY_NGAMR1 | 2015 | Nairobi | Male | >18 | Married | 1-2 | <0.016 | <0.002 | 0.064 | 0.064 | 0.38 | 0.25 | 8 |
| 2 | KNY_NGAMR2 | 2015 | Nairobi | Male | 26 | Single | 1-2 | <0.016 | <0.002 | 64 | 0.5 | 0.016 | 0.5 | 8 |
| 3 | KNY_NGAMR3 | 2015 | Nairobi | Male | 27 | Single | 1-2 | <0.016 | NT | 48 | 16 | 3 | 0.125 | 6 |
| 4 | KNY_NGAMR4 | 2016 | Coast | Male | 30 | Married | 1-2 | <0.016 | <0.002 | 8 | 12 | 0.006 | 0.125 | 2 |
| 5 | KNY_NGAMR5 | 2016 | Nyanza | Male | 18 | Single | 1-2 | <0.016 | 0.006 | 3 | 16 | 4 | 0.125 | 2 |
| 6 | KNY_NGAMR6 | 2017 | Coast | Male | 26 | Single | 1-2 | <0.016 | <0.016 | 0.094 | 0.75 | 12 | 0.25 | 3 |
| 7 | KNY_NGAMR7 | 2014 | Coast | Male | 19 | Single | 1-2 | <0.016 | 0.008 | >256 | 16 | 8 | 2 | 16 |
| 8 | KNY_NGAMR8 | 2013 | Nyanza | Female | 18 | Single | 1-2 | <0.016 | 0.002 | 0.064 | 0.5 | 3 | 0.25 | 4 |
| 9 | KNY_NGAMR9 | 2016 | Nyanza | Male | 23 | Single | 1-2 | <0.016 | <0.016 | 0.094 | 0.32 | 12 | 0.25 | 6 |
| 10 | KNY_NGAMR10 | 2016 | Nyanza | Male | 37 | Married | 1-2 | <0.016 | <0.016 | >256 | 32 | 8 | 0.38 | 16 |
| 11 | KNY_NGAMR11 | 2016 | Nyanza | Male | 31 | Single | 1-2 | <0.016 | <0.016 | 12 | 12 | 3 | 0.25 | 8 |
| 12 | KNY_NGAMR13 | 2014 | Rift Valley | Male | 25 | Married | 1-2 | <0.016 | <0.002 | 8 | 12 | 3 | 0.125 | 24 |
| 13 | KNY_NGAMR14 | 2015 | Rift Valley | Male | 28 | Married | 1-2 | <0.016 | <0.002 | 64 | 24 | 16 | 0.125 | 8 |
| 14 | KNY_NGAMR15 | 2014 | Nyanza | Male | 29 | Married | 1-2 | <0.016 | <0.016 | 48 | 32 | 4 | 0.5 | 12 |
| 15 | KNY_NGAMR16 | 2015 | Nyanza | Male | 30 | Married | 1-2 | <0.016 | <0.016 | 0.19 | 4 | 8 | 0.38 | 4 |
| 16 | KNY_NGAMR17 | 2015 | Nyanza | Male | 22 | Single | 9-10 | <0.016 | <0.016 | 0.38 | 16 | 6 | 0.5 | 6 |
| 17 | KNY_NGAMR18 | 2015 | Nyanza | Male | 22 | Single | None | <0.016 | <0.016 | 0.5 | 24 | 0.38 | 0.5 | 6 |
| 18 | KNY_NGAMR19 | 2015 | Nyanza | Male | 24 | Married | 1-2 | <0.016 | 0.094 | 0.19 | 64 | 4 | 1.5 | 1 |
| 19 | KNY_NGAMR20 | 2015 | Nyanza | Male | 34 | Married | 1-2 | <0.016 | 0.004 | 12 | 24 | 8 | 0.125 | 12 |
| 20 | KNY_NGAMR21 | 2016 | Nyanza | Male | 35 | Married | 1-2 | <0.016 | <0.016 | 32 | 24 | 3 | 0.125 | 12 |
| 21 | KNY_NGAMR22 | 2016 | Nyanza | Male | 22 | Single | 3-4 | <0.016 | 0.004 | 2 | 32 | 4 | 1 | 12 |
| 22 | KNY_NGAMR23 | 2017 | Nyanza | Female | 31 | Married | 1-2 | <0.016 | <0.002 | 12 | 8 | 12 | 0.125 | 4 |
| 23 | KNY_NGAMR24 | 2014 | Nyanza | Male | 27 | Single | 1-2 | <0.016 | <0.016 | 2 | 16 | 4 | 0.38 | 8 |
| 24 | KNY_NGAMR26 | 2016 | Nyanza | Male | 21 | Single | 1-2 | <0.016 | <0.016 | 8 | 32 | 1.5 | 0.25 | 8 |
| 25 | KNY_NGAMR28 | 2017 | Nyanza | Female | 24 | Married | 1-2 | <0.016 | <0.002 | 96 | 32 | 24 | 0.5 | 8 |
| 26 | KNY_NGAMR29 | 2017 | Nyanza | Male | 22 | Single | 1-2 | <0.016 | <0.002 | 0.094 | 12 | 12 | 0.25 | 2 |
| 27 | KNY_NGAMR30 | 2017 | Nyanza | Male | 19 | Single | 1-2 | <0.016 | <0.002 | 12 | 12 | 4 | 0.125 | 8 |
| 28 | KNY_NGAMR31 | 2017 | Nyanza | Male | 18 | Single | 1-2 | <0.016 | <0.002 | 0.38 | 48 | >32 | 0.25 | 3 |
| 29 | KNY_NGAMR32 | 2017 | Nyanza | Female | 26 | Married | 1-2 | <0.016 | <0.002 | 8 | 0.5 | 16 | 0.047 | 1.5 |
| 30 | KNY_NGAMR33 | 2016 | Nyanza | Male | 20 | Single | 1-2 | <0.016 | <0.016 | 3 | 32 | 2 | 0.25 | 4 |
| 31 | KNY_NGAMR35 | 2013 | Nairobi | Female | 26 | Married | 1-2 | NT | NT | >256 | 48 | 32 | 0.125 | NT |
| 32 | KNY_NGAMR41 | 2018 | Nyanza | Male | 42 | Single | 1-2 | <0.016 | <0.016 | 32 | 12 | 16 | NT | 4 |
| 33 | KNY_NGAMR50 | 2018 | Nairobi | Male | 26 | Single | 1-2 | <0.016 | <0.016 | 16 | 16 | 1.5 | 0.064 | 4 |
| 34 | KNY_NGAMR53 | 2018 | Rift Valley | Male | 35 | Married | 1-2 | <0.016 | <0.016 | 6 | 6 | 3 | 0.19 | 4 |
| 35 | KNY_NGAMR54 | 2018 | Rift Valley | Male | 35 | Married | 1-2 | <0.016 | <0.016 | 6 | 2 | 2 | 0.125 | 6 |

NT-Not tested, CFX- cefixime, CRO- ceftriaxone, PEN- penicillin, TET-tetracycline, CIP-ciprofloxacin, AZM-azithromycin.

**S2 Table.** EUCAST v8.0 (2018) Breakpoints for *N. gonorrhoeae*.

| **Antibiotic** | **Susceptible (S)** | **Resistant (R)** |
| --- | --- | --- |
| **Ceftriaxone (CRO)** | ≤ 0.125 mg/L | > 0.125 mg/L |
| **Cefixime (CFX)** | ≤ 0.125 mg/L | > 0.125 mg/L |
| **Azithromycin (AZM)** | ≤ 0.25 mg/L | > 0.5 mg/L |
| **Ciprofloxacin (CIP)** | ≤ 0.03 mg/L | > 0.06 mg/L |
| **Tetracycline (TET)** | ≤ 0.5 mg/L | > 1 mg/L |
| **Penicillin (PEN)** | ≤ 0.06 mg/L | > 1 mg/L |
| **Spectinomycin (SPT)** | ≤ 64 mg/L | > 64 mg/L |
| **Gentamicin (GEN)** | *No clinical breakpoints* | *No clinical breakpoints* |

**S3 Table.** Identified MLST alleles and sequence types. *Novel sequence types (STs) identified in this study are indicated in bold italics*

| Isolate | Region | Year of isolation | *abcZ* | *adk* | *aroE* | *fumC* | *gdh* | *pdhC* | *pgm* | MLST |
| --- | --- | --- | --- | --- | --- | --- | --- | --- | --- | --- |
| KNY_NGAMR1 | Nairobi | 2015 | 59 | 39 | 170 | 237 | 148 | 153 | 65 | **13613** |
| KNY_NGAMR2 | Nairobi | 2015 | 126 | 39 | 67 | 111 | 149 | 153 | 65 | 1928 |
| KNY_NGAMR3 | Nairobi | 2015 | 59 | 39 | 67 | 157 | 147 | 153 | 133 | 1893 |
| KNY_NGAMR4 | Coast | 2016 | 59 | 39 | 67 | 158 | 148 | 71 | 65 | 1588 |
| KNY_NGAMR5 | Nyanza | 2016 | 59 | 39 | 67 | 157 | 148 | 153 | 65 | 1599 |
| KNY_NGAMR6 | Coast | 2017 | 59 | 39 | 67 | 111 | 189 | 153 | 65 | 11367 |
| KNY_NGAMR7 | Coast | 2014 | 59 | 39 | 67 | 111 | 147 | 71 | 65 | 11366 |
| KNY_NGAMR8 | Nyanza | 2013 | 59 | 112 | 67 | 158 | 148 | 71 | 65 | 11365 |
| KNY_NGAMR9 | Nyanza | 2016 | 59 | 112 | 67 | 111 | 189 | 153 | 65 | **13614** |
| KNY_NGAMR10 | Nyanza | 2016 | 59 | 39 | 67 | 78 | 189 | 153 | 65 | 1932 |
| KNY_NGAMR11 | Nyanza | 2014 | 59 | 112 | 67 | 158 | 148 | 71 | 65 | 11365 |
| KNY_NGAMR13 | Rift Valley | 2015 | 59 | 112 | 937 | 158 | 148 | 71 | 65 | **13782** |
| KNY_NGAMR14 | Rift Valley | 2014 | 59 | 39 | 170 | 78 | 189 | 153 | 65 | 11242 |
| KNY_NGAMR15 | Nyanza | 2015 | 59 | 112 | 937 | 158 | 148 | 71 | 65 | **13782** |
| KNY_NGAMR16 | Nyanza | 2015 | 109 | 39 | 67 | 157 | 150 | 153 | 133 | **13766** |
| KNY_NGAMR17 | Nyanza | 2015 | 59 | 39 | 67 | 111 | 148 | 71 | 65 | 1921 |
| KNY_NGAMR18 | Nyanza | 2015 | 59 | 39 | 170 | 158 | 148 | 71 | 65 | 8133 |
| KNY_NGAMR19 | Nyanza | 2015 | 59 | 39 | 67 | 157 | 147 | 153 | 133 | 1893 |
| KNY_NGAMR20 | Nyanza | 2016 | 59 | 112 | 67 | 158 | 147 | 71 | 65 | **13780** |
| KNY_NGAMR21 | Nyanza | 2016 | 59 | 112 | 67 | 158 | 148 | 153 | 65 | 11976 |
| KNY_NGAMR22 | Nyanza | 2017 | 59 | 39 | 170 | 158 | 148 | 71 | 65 | 8133 |
| KNY_NGAMR23 | Nyanza | 2014 | 59 | 112 | 67 | 158 | 147 | 71 | 65 | **13780** |
| KNY_NGAMR24 | Nyanza | 2016 | 59 | 39 | 170 | 158 | 148 | 71 | 65 | 8133 |
| KNY_NGAMR26 | Nyanza | 2016 | 109 | 39 | 67 | 983 | 150 | 153 | 65 | **13779** |
| KNY_NGAMR28 | Nyanza | 2017 | 59 | 39 | 67 | 78 | 189 | 153 | 65 | 1932 |
| KNY_NGAMR29 | Nyanza | 2017 | 109 | 39 | 67 | 157 | 150 | 153 | 65 | 11750 |
| KNY_NGAMR30 | Nyanza | 2017 | 59 | 112 | 67 | 157 | 148 | 71 | 65 | **13763** |
| KNY_NGAMR31 | Nyanza | 2017 | 59 | 39 | 67 | 78 | 189 | 153 | 65 | 1932 |
| KNY_NGAMR32 | Nyanza | 2017 | 59 | 112 | 937 | 158 | 148 | 71 | 65 | **13782** |
| KNY_NGAMR33 | Nyanza | 2016 | 109 | 39 | 67 | 157 | 150 | 901 | 133 | **13764** |
| KNY_NGAMR35 | Nairobi | 2013 | 59 | 39 | 67 | 78 | 189 | 153 | 65 | 1932 |
| KNY_NGAMR41 | Nyanza | 2018 | 59 | 39 | 170 | 158 | 148 | 71 | 65 | 8133 |
| KNY_NGAMR50 | Nairobi | 2018 | 59 | 39 | 67 | 78 | 189 | 153 | 65 | 1932 |
| KNY_NGAMR53 | Rift Valley | 2018 | 109 | 39 | 67 | 157 | 150 | 153 | 65 | 11750 |
| KNY_NGAMR54 | Rift Valley | 2018 | 126 | 39 | 170 | 158 | 148 | 71 | 65 | 8111 |

**S4 Table.**
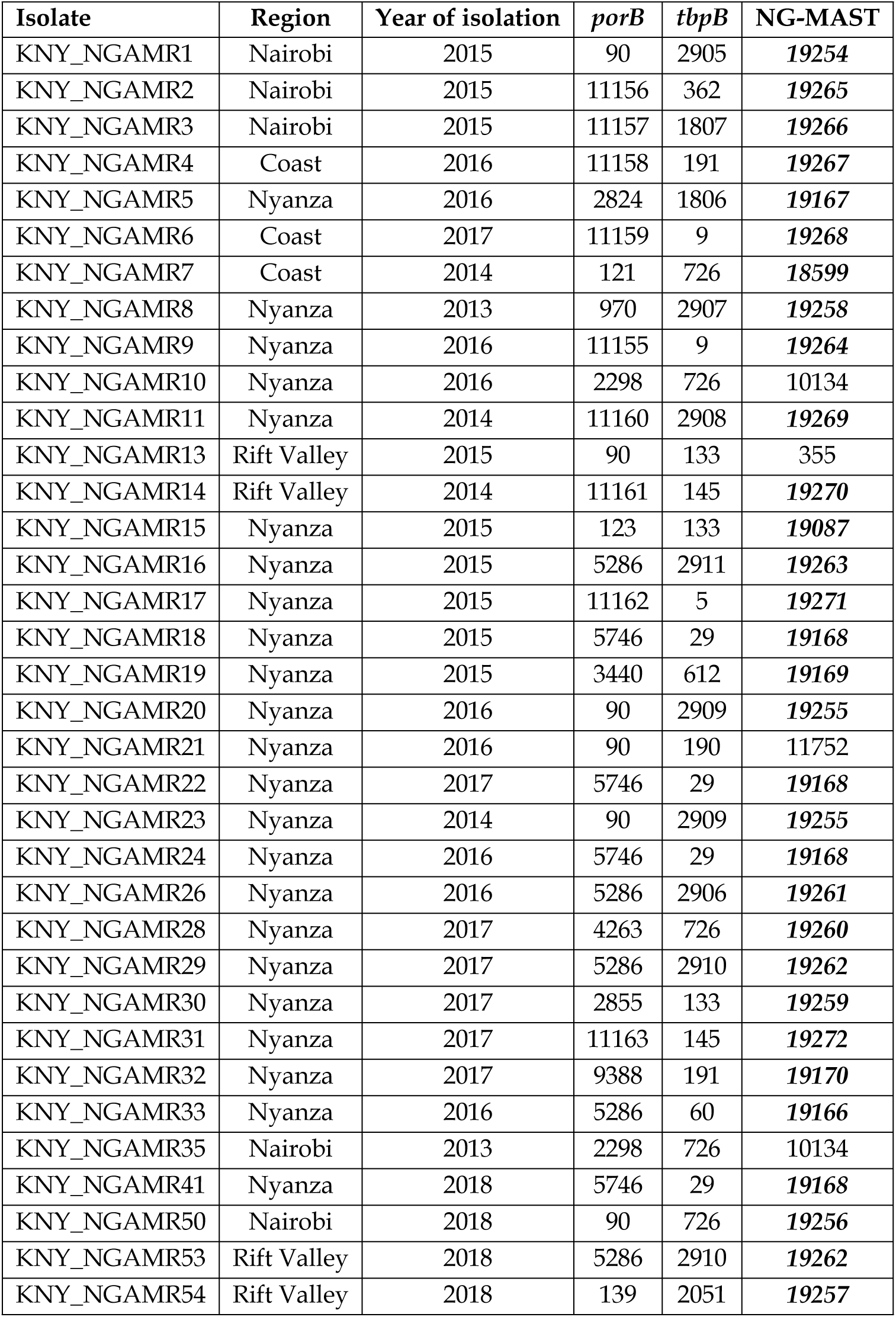
Identified NG-MAST alleles and sequence types. *Novel sequence types (STs) identified in this study are indicated in bold italics*

**S5 Table.** Identified NG-STAR alleles and sequence types. Novel sequence types (STs) identified in this study are indicated in bold italics

| Isolate | Region | *mtrR* | *penA* | 23S_rRNA | *gyrA* | *parC* | *ponA* | *porB* | NG STAR |
| --- | --- | --- | --- | --- | --- | --- | --- | --- | --- |
| KNY_NGAMR1 | Nairobi | 57 | 294 | 100 | 1 | 2 | 100 | 13 | 2661 |
| KNY_NGAMR2 | Nairobi | 19 | 23 | 100 | 100 | 2 | 1 | 4 | 2050 |
| KNY_NGAMR3 | Nairobi | 10 | 285 | 100 | 1 | 49 | 1 | 100 | 1054 |
| KNY_NGAMR4 | Coast | 54 | 228 | 100 | 100 | 1 | 100 | 3 | 2664 |
| KNY_NGAMR5 | Nyanza | 229 | 285 | 100 | 1 | 22 | 1 | 19 | 2665 |
| KNY_NGAMR6 | Coast | 10 | 23 | 100 | 1 | 22 | 100 | 100 | 1271 |
| KNY_NGAMR7 | Coast | 98 | 20 | 100 | 1 | 49 | 1 | 3 | 1516 |
| KNY_NGAMR8 | Nyanza | 10 | 294 | 100 | 7 | 7 | 1 | 13 | 1586 |
| KNY_NGAMR9 | Nyanza | 10 | 294 | 100 | 7 | 22 | 100 | 14 | 2660 |
| KNY_NGAMR10 | Nyanza | 343 | 23 | 100 | 1 | 22 | 100 | 100 | **3186** |
| KNY_NGAMR11 | Nyanza | 10 | 294 | 100 | 7 | 7 | 1 | 13 | 1586 |
| KNY_NGAMR13 | Rift Valley | 10 | 294 | 100 | 7 | 7 | 1 | 13 | 1586 |
| KNY_NGAMR14 | Rift Valley | 10 | 23 | 100 | 1 | 22 | 100 | 3 | 1255 |
| KNY_NGAMR15 | Nyanza | 10 | 294 | 100 | 7 | 7 | 1 | 14 | 2170 |
| KNY_NGAMR16 | Nyanza | 10 | 294 | 100 | 1 | 22 | 100 | 14 | 1890 |
| KNY_NGAMR17 | Nyanza | 18 | 20 | 100 | 1 | 49 | 1 | 100 | 2667 |
| KNY_NGAMR18 | Nyanza | 54 | 228 | 100 | 1 | 49 | 100 | 100 | 2668 |
| KNY_NGAMR19 | Nyanza | 230 | 294 | 100 | 1 | 49 | 1 | 100 | 2669 |
| KNY_NGAMR20 | Nyanza | 10 | 285 | 100 | 7 | 49 | 1 | 13 | 2666 |
| KNY_NGAMR21 | Nyanza | 10 | 294 | 100 | 7 | 7 | 1 | 13 | 1586 |
| KNY_NGAMR22 | Nyanza | 54 | 228 | 100 | 1 | 49 | 100 | 100 | 2668 |
| KNY_NGAMR23 | Nyanza | 10 | 285 | 100 | 7 | 49 | 1 | 13 | 2666 |
| KNY_NGAMR24 | Nyanza | 54 | 228 | 100 | 1 | 49 | 100 | 100 | 2668 |
| KNY_NGAMR26 | Nyanza | 10 | 294 | 100 | 1 | 22 | 100 | 14 | 1890 |
| KNY_NGAMR28 | Nyanza | 10 | 23 | 100 | 7 | 22 | 100 | 14 | **3184** |
| KNY_NGAMR29 | Nyanza | 10 | 294 | 100 | 1 | 111 | 1 | 14 | **3179** |
| KNY_NGAMR30 | Nyanza | 10 | 294 | 100 | 7 | 7 | 1 | 14 | 2170 |
| KNY_NGAMR31 | Nyanza | 10 | 23 | 100 | 1 | 22 | 100 | 13 | **3183** |
| KNY_NGAMR32 | Nyanza | 10 | 294 | 100 | 7 | 22 | 1 | 13 | 1603 |
| KNY_NGAMR33 | Nyanza | 10 | 294 | 100 | 1 | 22 | 100 | 14 | 1890 |
| KNY_NGAMR35 | Nairobi | 10 | 23 | 100 | 1 | 22 | 100 | 100 | 1271 |
| KNY_NGAMR41 | Nyanza | 54 | 228 | 100 | 1 | 49 | 100 | 100 | 2668 |
| KNY_NGAMR50 | Nairobi | 10 | 23 | 100 | 7 | 22 | 100 | 13 | **3185** |
| KNY_NGAMR53 | Rift Valley | 10 | 294 | 100 | 1 | 111 | 1 | 14 | **3179** |
| KNY_NGAMR54 | Rift Valley | 10 | 20 | 100 | 1 | 22 | 100 | 100 | **3182** |

**S6 Table.**
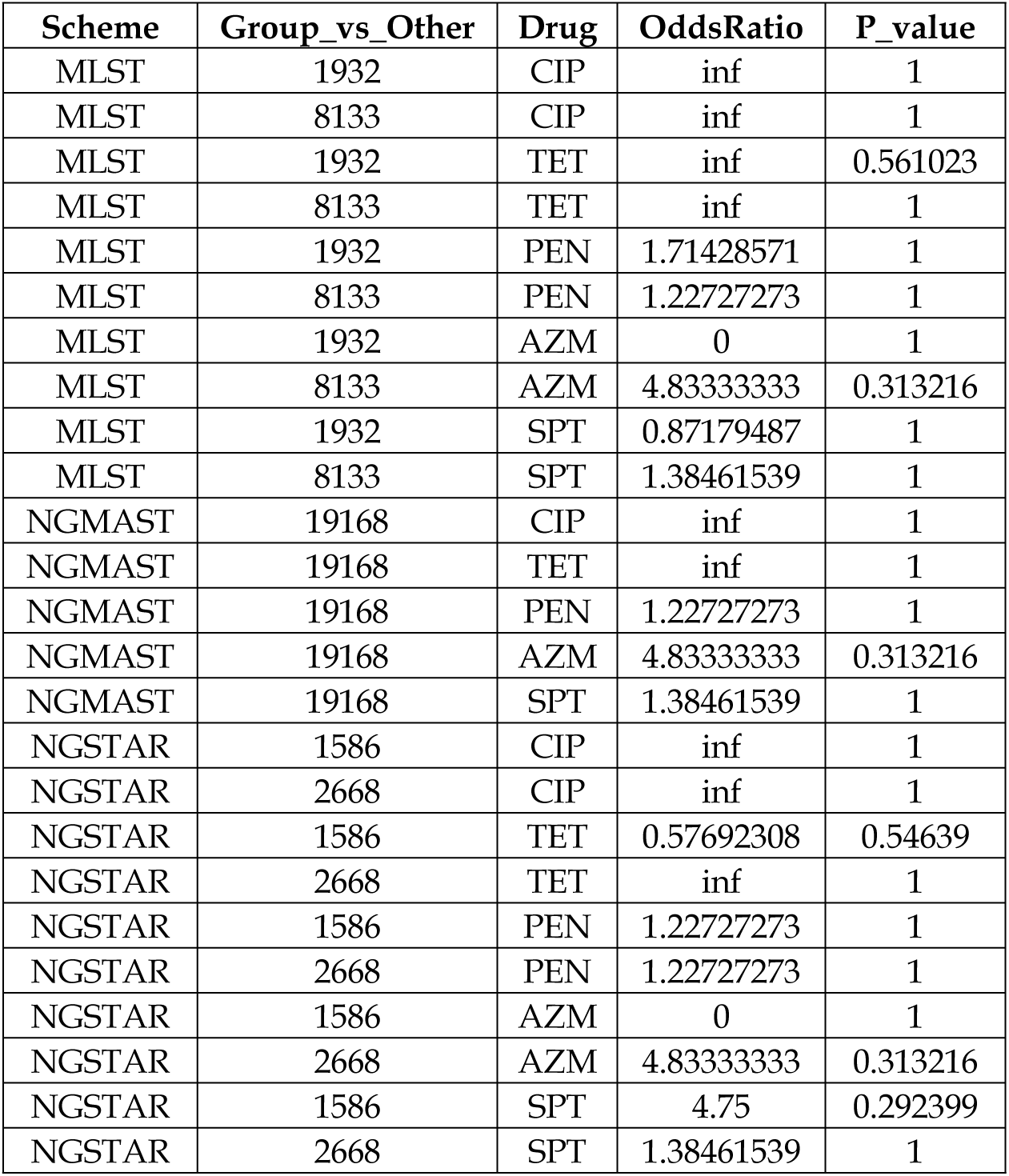
Typing scheme and AMR associations.

| **Scheme** | **Group_vs_Other** | **Drug** | **OddsRatio** | **P_value** |
| --- | --- | --- | --- | --- |
| MLST | 1932 | CIP | inf | 1 |
| MLST | 8133 | CIP | inf | 1 |
| MLST | 1932 | TET | inf | 0.561023 |
| MLST | 8133 | TET | inf | 1 |
| MLST | 1932 | PEN | 1.71428571 | 1 |
| MLST | 8133 | PEN | 1.22727273 | 1 |
| MLST | 1932 | AZM | 0 | 1 |
| MLST | 8133 | AZM | 4.83333333 | 0.313216 |
| MLST | 1932 | SPT | 0.87179487 | 1 |
| MLST | 8133 | SPT | 1.38461539 | 1 |
| NGMAST | 19168 | CIP | inf | 1 |
| NGMAST | 19168 | TET | inf | 1 |
| NGMAST | 19168 | PEN | 1.22727273 | 1 |
| NGMAST | 19168 | AZM | 4.83333333 | 0.313216 |
| NGMAST | 19168 | SPT | 1.38461539 | 1 |
| NGSTAR | 1586 | CIP | inf | 1 |
| NGSTAR | 2668 | CIP | inf | 1 |
| NGSTAR | 1586 | TET | 0.57692308 | 0.54639 |
| NGSTAR | 2668 | TET | inf | 1 |
| NGSTAR | 1586 | PEN | 1.22727273 | 1 |
| NGSTAR | 2668 | PEN | 1.22727273 | 1 |
| NGSTAR | 1586 | AZM | 0 | 1 |
| NGSTAR | 2668 | AZM | 4.83333333 | 0.313216 |
| NGSTAR | 1586 | SPT | 4.75 | 0.292399 |
| NGSTAR | 2668 | SPT | 1.38461539 | 1 |

## Notes

### Competing Interest Statement

The authors have declared no competing interest.

